# Human shields alter antipredator behavior in Guenther’s dik-dik (*Madoqua guentheri*)

**DOI:** 10.1101/2025.09.07.674752

**Authors:** Raymond O. Owino, Isobel Hawkins, Iris Noordermeer, Margaret Mercer, Lauren A. Stanton, Jacob R. Goheen, Jesse M. Alston

**Affiliations:** University of Arizona, School of Natural Resources, and the Environment, 1064 E. Lowell St, Tucson, Arizona, USA 85721; University of Oxford, Department of Biology, 11a Mansfield Rd, Oxford OX1 3SZ, UK; Leiden University, Institute of Biology, Sylviusweg 72, 2333BE Leiden, The Netherlands; University of California Berkeley, Department of Environmental Science, Policy, and Management, 130 Mulford Hall, Berkeley, CA, USA 94720; Iowa State University, Department of Natural Resource Ecology and Management, 2310 Pammel Dr, Ames, Iowa, USA; Mpala Research Centre, Rumuruti Road, Nanyuki, Kenya

## Abstract

Human activities affect landscape use by wildlife, with many predators actively avoiding areas near humans. In human-occupied areas, prolonged habituation to human activity could therefore lead to relaxed antipredator behavior by prey species, which can be detrimental when predators return or if individuals are translocated to new areas that host predators. We conducted behavioral trials to test whether exposure of Guenther’s dik-dik (*Madoqua guenther*i) to human activity affected habituation to humans and, by extension, response to predation cues. We hypothesized that dik-diks living in areas with higher levels of human activity would exhibit shorter flight initiation distances and spend less time responding to predation cues than those in areas with less human activity, but would respond more strongly to non-specific predation cues (i.e., alarm calls from white-browed sparrow-weavers, a locally common bird) than to cues associated with predators that avoid human activity (i.e., hyena vocalizations). Flight initiation distance and responses to both sparrow-weaver alarm calls and hyena vocalizations varied predictably with differences in human activity: dik-diks living in areas with more human activity had shorter flight initiation distances and spent less time responding to predation cues than those living in areas with less human activity, but responded more strongly to the sparrow-weaver alarm calls than hyena vocalizations in the area with the highest level of human activity. As human populations expand and overlap increasingly with predators, human settlements and activities may increase susceptibility of prey to predators by increasing prey naivety to predators that avoid humans. These results are particularly relevant for ecotourism in working landscapes and translocation of habituated animals, both of which may increase prey naivety.

## 1. Introduction

Globally, the extent of landscapes free of human presence and activity is decreasing (Corlett, 2015; Mu et al., 2022) leading to more frequent interactions between humans and other species (Ellis, 2011; Worm & Paine, 2016; Moll et al., 2021). When this happens, fear of humans as apex predators can have a cascading effect on landscape use by wildlife, constraining where wildlife occurs within landscapes (Laundré et al., 2001a; Smith et al., 2017; Suraci et al., 2019). Even where humans are not actively controlling wildlife lethally (through hunting, poaching, or persecution), human activity within landscapes disproportionally affects large predators (Woodroffe & Frank, 2005), with many strongly avoiding areas near humans (Kautz et al., 2021; Barker et al., 2023). Because large predators in particular tend to avoid human activity (Laundré et al., 2001b; Muhly et al., 2011; Suraci et al., 2019), humans can create areas in which prey species experience reduced risk of predation from large predators (often called the “human shield hypothesis”; Berger, 2007).

Wildlife are naturally afraid of humans and in most cases respond by fleeing from them (Knight, 2009). Upon initial exposure to humans, time and energy spent running as opposed to feeding and breeding could be detrimental (Arlettaz et al., 2015). Where humans do not hunt or poach wildlife, avoidance of humans could impose opportunity costs (e.g., reduced foraging and reproduction; Pérez-Tris et al., 2004; Ciuti et al., 2012; Smith et al., 2017). Where vigilance and other costly antipredator behaviors are no longer necessary, reallocating time and energy to feeding and breeding should increase fitness (Blumstein & Daniel, 2005). As a result, where wildlife experience prolonged exposure to benign interactions with humans, it can lead to habituation (i.e., loss of response to stimuli that have little or no effect on survival; Bejder et al., 2009; Blumstein, 2016).

Habitation to humans allows prey species to exploit human shields (Pérez-Flores et al., 2022). In human-occupied areas, however, release from the risk of predation (predator absence) may lead to naivety to nonhuman predators (Geffroy et al., 2015; Blumstein, 2016; D’Ammando & Bro-Jørgensen, 2023; Wheat & Wilmers, 2016). While some antipredator behavior may be instinctual (and thus fixed), learning allows individuals to fine-tune antipredator behavior to local conditions (Griffin, 2004; Crane & Ferrari, 2013; Wooster et al., 2024). Where prey species are protected from predators, antipredator behaviors that are costly (e.g., vigilance, habitat use) are expected to become less common (Blumstein & Daniel, 2005; Creel, 2018; Saltz et al., 2019). Since individual survival depends on timely responses to predation cues, relaxed antipredator behavior could be costly should predation risk increase via predator introduction or reintroduction (Berger, 1999; Ford et al., 2015).

The number of predator species decreases as human activity increases (Muhly et al., 2011), but predator avoidance of humans is largely based on behavioral conditioning (Zinn et al., 2008). Since predator species vary in the degree to which they avoid humans, human shields are not uniformly effective against all predator species (Van Scoyoc et al., 2023), resulting in prey naivety only to those predators excluded by human shields. So, while cues that are specific to a predator species (e.g., vocalizations by a human shield-excluded predator like hyenas) may be ignored by prey that are naive to them, non-specific predation cues (e.g., alarm calls from other species) could still elicit antipredator behavior (Meise et al., 2018; Turner et al., 2023). Testing responses of prey species to both specific and non-specific predator cues may therefore reveal the extent to which human activity results in relaxed antipredator behavior specific to only those predators excluded by human activity, or to all predators more generally.

We tested whether human activity affects habituation to humans, and by extension, the relaxation of antipredator responses of Guenther’s dik-dik (*Madoqua guentheri*) to predator-specific and non-specific predation cues. Because prolonged exposure to humans can lead to habituation to humans, we hypothesized that dik-diks living in areas of higher human activity respond less strongly to human presence than those living in areas of lower human activity (Hypothesis 1). Because large predators tend to avoid areas with high human activity (Suraci et al., 2019; Kautz et al., 2021; Barker et al., 2023), and release from predation pressure leads to predator naivety in prey species when isolated from predators (Berger, 1999; Berger et al., 2001; Blumstein & Daniel, 2005), we hypothesized that dik-diks in areas of higher human activity would respond less strongly to both predator-specific (Hypothesis 2) and non-specific predation cues (Hypothesis 3) than dik-diks living in areas of lower human activity. Finally, we expected that dik-diks occurring in areas of high human activity would respond more strongly to non-specific predation cues than to cues from predators typically excluded by human activity (Hypothesis 4).

## 2. Methods

### 2.1. Study area and species

We conducted our experiment within Mpala Research Centre and Conservancy (hereafter “Mpala”; 0° 17′ N, 37° 53′ E,1600 m elevation) in Laikipia County in central Kenya. Mpala is a privately owned property that covers about 194 km^2^, bordered by the Ewaso Ng’iro (River of Brown Waters) along the northern and eastern side. The area is generally characterized by semi-arid savannas dominated by woody *Acacia* species (*Acacia* [*Senegalia*] *drepanolobium*, *A.* [*Vachellia*] *brevispica, A.* [*V.*] *etbaica*, *A.* [*S.*] *mellifera*) and grasses (primarily *Themeda, Cynodon* and *Pennisetum* species). Mpala is located in the rain shadow of Mount Kenya, which imposes pronounced climatic variability at relatively small spatial scales: from 2009–2019, mean annual rainfall increased >15% across the property from north to south, and historically this gradient was even stronger. Our study site was located on the southern portion of Mpala, with mean rainfall ranging from 369 to 839 mm per year, and temperatures ranging from 12.4°C to 28.1°C on a daily basis and varying little across the year (Alston et al., 2022).

Because hunting is prohibited in Kenya and because security patrols minimize poaching, free-ranging wildlife often encounter and thus habituate to humans. Wild ungulates on Mpala include plains and Grevy’s zebra (*Equus quagga* and *E. grevyi*, respectively), reticulated giraffe (*Giraffa reticulata*), African buffalo (*Syncerus caffer*), impala (*Aepyceros melampus*), African bush elephant (*Loxodonta africana*) and Guenther’s dik-dik (hereafter “dik-diks”; Augustine, 2010; Alston et al., 2022). The most common native large predators are lions (*Panthera leo*), leopards (*Panthera pardus*), spotted hyenas (*Crocuta crocuta*), and African wild dogs (*Lycaon pictus*). Because these large predators often face retaliation after preying on livestock, they are more sensitive to human activities than the herbivores (Ikanda & Packer, 2008; Ontiri et al., 2019).

Dik-diks are among the smallest ungulates and thus are highly vulnerable to predation from a wide array of predators at Mpala, including leopards, spotted hyenas, African wild dogs, silver-backed jackals (*Canis mesomelas*), wildcats (*Felis silvestris*), caracals (*Caracal caracal*), servals (*Leptailurus serval*), cheetahs (*Acinonyx jubatus*), olive baboons (*Papio anubis*), Verreaux’s eagles (*Aquila verreauxii*), and martial eagles (*Polemaetus bellicosus*; Kingswood & Kumamoto, 1996; Augustine, 2010). They therefore spend a substantial amount of time in a vigilant state, sometimes eavesdropping on alarm calls from unrelated species such as the white-bellied go-away bird (*Corythaixoides leucogaster*; Lea et al., 2008). Dik-diks are relatively sedentary and they are faithful to small (<1 ha) territories, even when exposed to predator scent (Ford & Goheen, 2015). Therefore, in human-dominated landscapes, they may experience chronically high levels of human activities without abandoning their territories, which could lead to habituation to humans thus allowing them to exploit human shields.

### 2.2. Behavioral trials

We stratified our study area into three sites according to levels of human activity (Fig. 1). To represent relatively low levels of human activity, we selected a section of private road along the Ewaso Ng’iro traversed only by vehicle and with relatively little traffic. To represent areas of moderate human activity, we selected an intermittently inhabited camp along the Ewaso Ng’iro that is used for field courses and overflow accommodation. For an area of high human activity, we selected the main research center, characterized by permanent settlement with high levels of human activity year-round.

**Figure 1:**
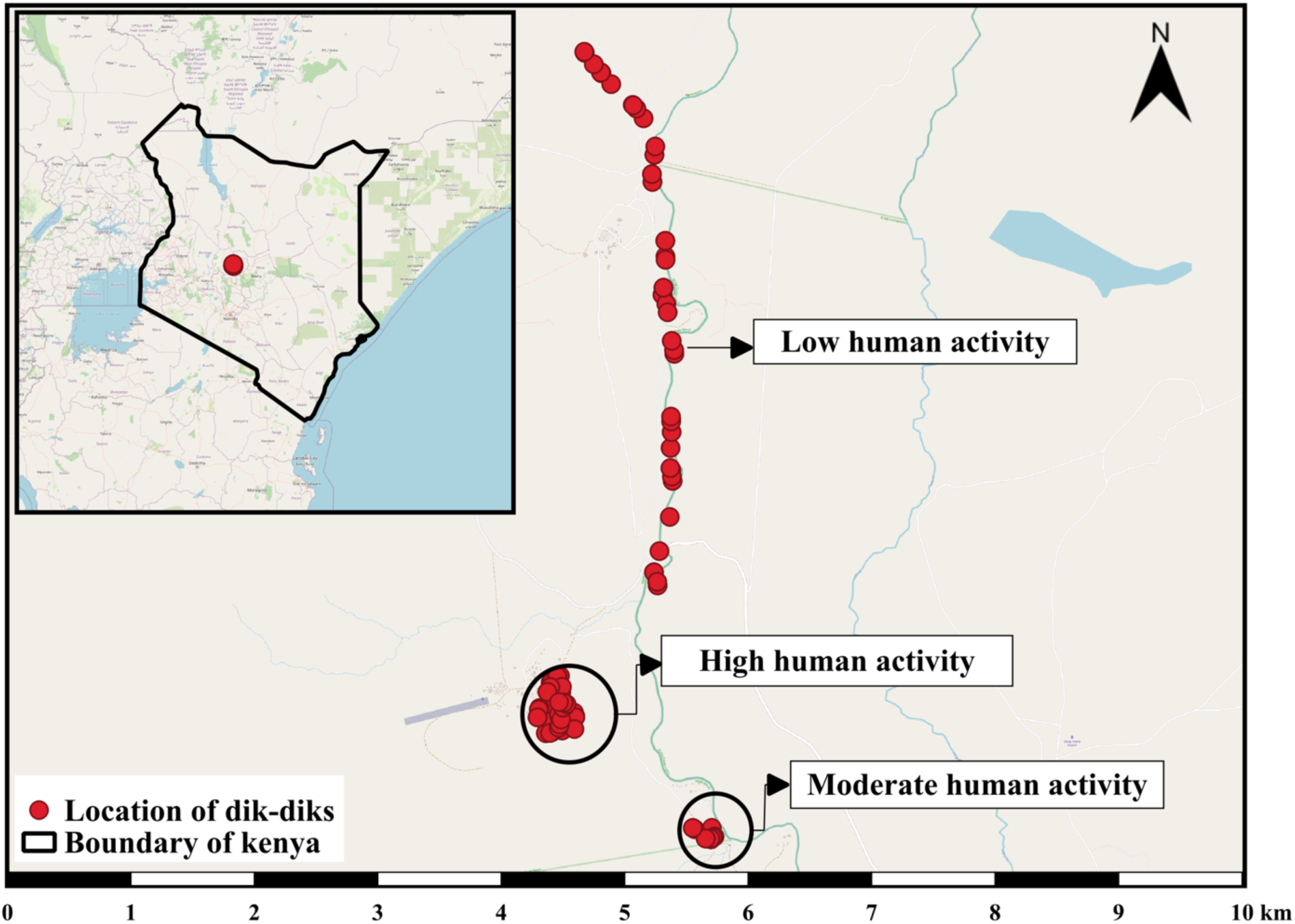
Map of study site. Insert map represents Kenya’s boundary with red dot showing the location of Mpala Research Centre and Conservancy. Red dots in main map represent dik-dik locations across the human activity gradient (from top: low, high, and moderate human activity, respectively). Basemap is OpenStreetMap (OSM) standard from QuickMapServices plugin.

We observed dik-diks in August and September 2022 during daylight hours (0600 hrs to 1800 hrs). We used flight initiation distance (i.e., the distance between an approaching human and an animal when the animal begins to flee; Weston et al., 2012) to compare how variation in exposure to human activity influenced habituation to humans by dik-diks. We interpreted shorter flight initiation distances as indicative of habituation to humans (Engelhardt & Weladji, 2011; Bateman & Fleming, 2014; Mbise et al., 2020). Within each site, the same observer approached dik-diks at a rate of approximately one step per second. Then, using a laser rangefinder (Nikon Prostaff 1000, Tokyo, Japan), we measured the distance between the observer and the dik-dik’s final location before it fled the approaching observer. We also recorded coordinates for the location of the observer and the bearing of the dik-dik’s location. Occasionally, and especially in areas of low human activity, dik-diks fled before we could approach them. In this case, we recorded the distance from where we stood to the location at which the dik-dik was first spotted fleeing.

Increased human activity has been linked to decreased antipredator behavior, with individuals farther from centers of human activity exhibiting heightened responses to predation cues and allocating more time to antipredator behavior (Berger, 1999; Coleman et al., 2008). To compare how spatial variation in human activity affected the amount of time dik-diks spend on antipredator behavior, we exposed dik-diks to two types of predation cue and a control treatment through playback recordings. We used playback recordings that represented a predator-specific predation cue, a non-specific predation cue, and a nonthreatening control treatment: (1) spotted hyena (*Crocuta crocuta*) vocalizations, (2) white-browed sparrow-weaver (*Plocepasser mahali*) alarm calls, and (3) sooty boubou (*Laniarius leucorhynchus*) songs, respectively. Spotted hyenas are a common predator of dik-dik at Mpala and often calls at night at all three study sites. They avoid human activity, particularly during the day. White-browed sparrow-weavers are common cooperative breeders that feed in groups. One individual in the group remains on guard to make alarm calls when predators are present (Ferguson, 1987; Voigt et al., 2021). The sooty boubou is a common songbird at Mpala that sings frequently, but that does not emit alarm calls. We obtained three recordings of spotted hyenas feeding at carcasses from a variety of sources including YouTube and a local ranger (Call #1 [Lion vs hyena epic audio, MalaMala Game Reserve, YouTube], Call #2 [Hyena spotted hyena in the road calling, 100% Nature, YouTube], and Call #3, [S. Haji, personal communication]). Using a hand-held recorder (Olympus LS-12, Tokyo, Japan) and a shotgun microphone (Sennheiser ME-66, Wedemark, German), we recorded sparrow-weaver alarm calls in response to a kite shaped like a hawk flown at three different nest sites, as well as boubous singing at three different sites. To reduce noise and improve the quality of audio recordings, we edited each sound clip, trimmed it to twenty seconds, and normalized the sound to 75 decibels (dB) using SASlab Pro (Avisoft, Brandenburg, Germany). We then located dik-diks by walking on planned routes and broadcasted the recordings through a wireless Bluetooth speaker (Ultimate Ears, California, United States) for 20 seconds. Within each site, the same recorder recorded dik-dik behavioral changes over an 80-second observation period (including 20 seconds during the broadcasted calls) using an ethogram (Table 1) modified from Lea et al. (2008). It is important to note dik-diks could be engaged in any of these activities simultaneously, such as sitting while being vigilant. We therefore assumed there was no response when dik-dik continued an activity in which it already was engaged. To reduce the risk of nearby dik-diks overhearing the playback ahead of our approach, we rotated systematically between predation cues (hyena vocalizations or sparrow-weaver alarm calls) and the control treatment (boubou song). We predominantly broadcasted the calls while on foot, except in the low human activity site, where dik-diks often fled before we could place the speaker on the ground. To work around this, we used a vehicle to approach dik-diks, then halted the engine and allowed time for dik-diks to return to normal activities (Table 1) before broadcasting the recording through the window.

**Table 1:** Modified ethogram of (Lea et al., 2008) to score behavioral transitions when playing playback sounds.

| Response Category | Type of activity |
| --- | --- |
| No response | Calm<br>Grooming<br>Scratching<br>Sitting<br>Foraging<br>Ruminating |
| Vigilance | Standing<br>Nose Twitching<br>Tongue Flicking<br>Looking |
| Flight | Walking from observer<br>Running from observer |

### 2.3. Statistical analysis

All statistical analyses were performed using the R statistical software environment (v4.4.0; R Core Team, 2023). To explore how human activity affected flight initiation distances by dik-diks along the human activity gradient, we carried out a simple linear regression with the three sites as a categorical explanatory variable, equivalent to a one-way Analysis of Variance (ANOVA). For ease of interpretation, we used the low human activity site as the reference category.

To quantify how human activity affected the proportion of time spent by dik-diks in response to predation cues, we analyzed the proportion of time dik-diks spent exhibiting different responses: normal activity was considered as a state of “no response”, while vigilance and flight were summed to represent a state of “response” to the broadcasted calls (Table 1). It is important to note that this proportion has a continuous distribution commensurate to beta distribution but can include 0 and 1. The data therefore violates the assumptions of beta regression since there is some probability of observing the same exact value multiple times (particularly 0s and 1s). The proportion could also be characterized with a binomial distribution, though it does not quite represent traditional 0s and 1s. We therefore chose to analyze our behavioral trial data using binomial-family generalized linear models because this approach is more common across wildlife biology than beta regression and therefore likely familiar to more readers. Results of beta regression are provided in Appendix S1. To quantify the effect of human activity on dik-dik responses, we first tested how dik-diks responded to the three types of calls broadcast to represent different types of predation cues. This was to confirm that dik-diks can distinguish the difference between “threatening” (i.e., hyena vocalizations, sparrow-weaver alarm calls) and “nonthreatening” (i.e., boubou song) sounds. For the three sites, we performed separate generalized linear models (binomial family, logit link) to quantify the association between the response variable (proportion of time dik-diks spent responding to calls) and the type of predation cue (predator-specific, non-specific, and control) as the explanatory variable. Because we did not expect dik-diks to respond to the control treatment (boubou song; Lea et al., 2008), we used this as the reference category.

Finally, we tested whether responses to types of predation cue (predator-specific, non-specific, and control) varied along the human activity gradient. For the three types of predation cue, we performed another series of generalized linear models (binomial family, logit link) to quantify the association between the response variable (proportion of time dik-diks spent responding to predation cues) and the human activity gradient (low, moderate, and high) as the explanatory variable. We used our site with low human activity as our reference category.

## 3. Results

To examine the influence of human activity on flight initiation distance, we conducted 101 behavioral trials along the three sites (20, 38, and 43 trials in areas of low, moderate, and high human activity, respectively). Flight initiation distances for dik-diks decreased with increasing human activity (Fig. 2). Dik-diks in areas of low human activity had the longest flight initiation distances, 45.8 m (95% CI: 37.0 – 54.5 m), averaging 18.8 m (95% CI: 8.0 – 29.6 m; *p* < 0.001), and 38.5 m (95% CI: 28.0 – 49.1 m; *p* < 0.001) farther than flight initiation distances for dik-diks in areas of moderate and high human activity, respectively.

**Figure 2:**
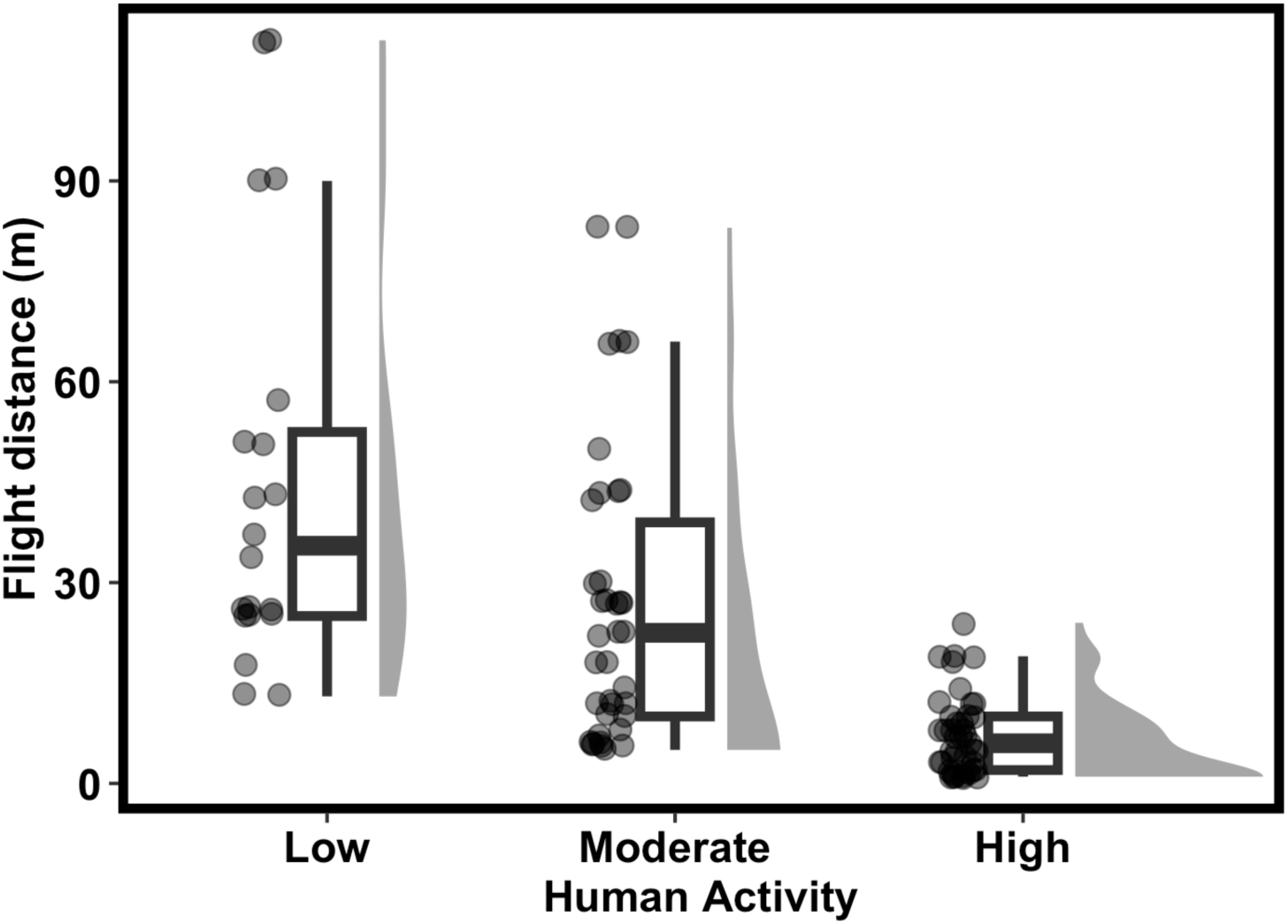
Flight initiation distances across the human activity gradient. Overall flight initiation distances decreased with increases in human activity. Box-and-whisker components indicate quantiles of data, points on the left of each treatment represent individual data points, and the distributions on the right of each treatment represent frequency distributions of the data.

To examine the influence of human activity on antipredator behavior by dik-diks, we conducted 122 behavioral trials along the three sites (49, 15, and 58 trials in areas of low, moderate, and high human activity, respectively). Overall, dik-diks responded more strongly to both hyena vocalizations and sparrow-weaver alarm calls than sooty boubou song (Fig. 3). In areas of low human activity (Fig. 3A), the odds of dik-diks responding increased by 284% (95% CI: 227 – 351%; *p* < 0.001) and 147% (95% CI: 110 – 189%; *p* < 0.001) to hyena vocalizations and sparrow-weaver alarm calls, respectively, compared to the sooty boubou song. In areas of moderate human activity (Fig. 3B), the odds of dik-diks responding increased by 1845% (95% CI: 1172 – 2997%; *p* < 0.001) and 776% (95% CI: 460 – 1321%; *p* < 0.001) when responding to hyena vocalizations and sparrow-weaver alarm calls, respectively, compared to the sooty boubou song. In areas of high human activity (Fig. 3C), the odds of dik-diks responding did not differ (95% CI: -20 – 20%; *p* = 0.833) when responding to hyena vocalizations compared to sooty boubou song, but increased by 137% (95% CI: 95 – 191%; *p* < 0.001) when responding to sparrow-weaver alarm calls.

**Figure 3:**
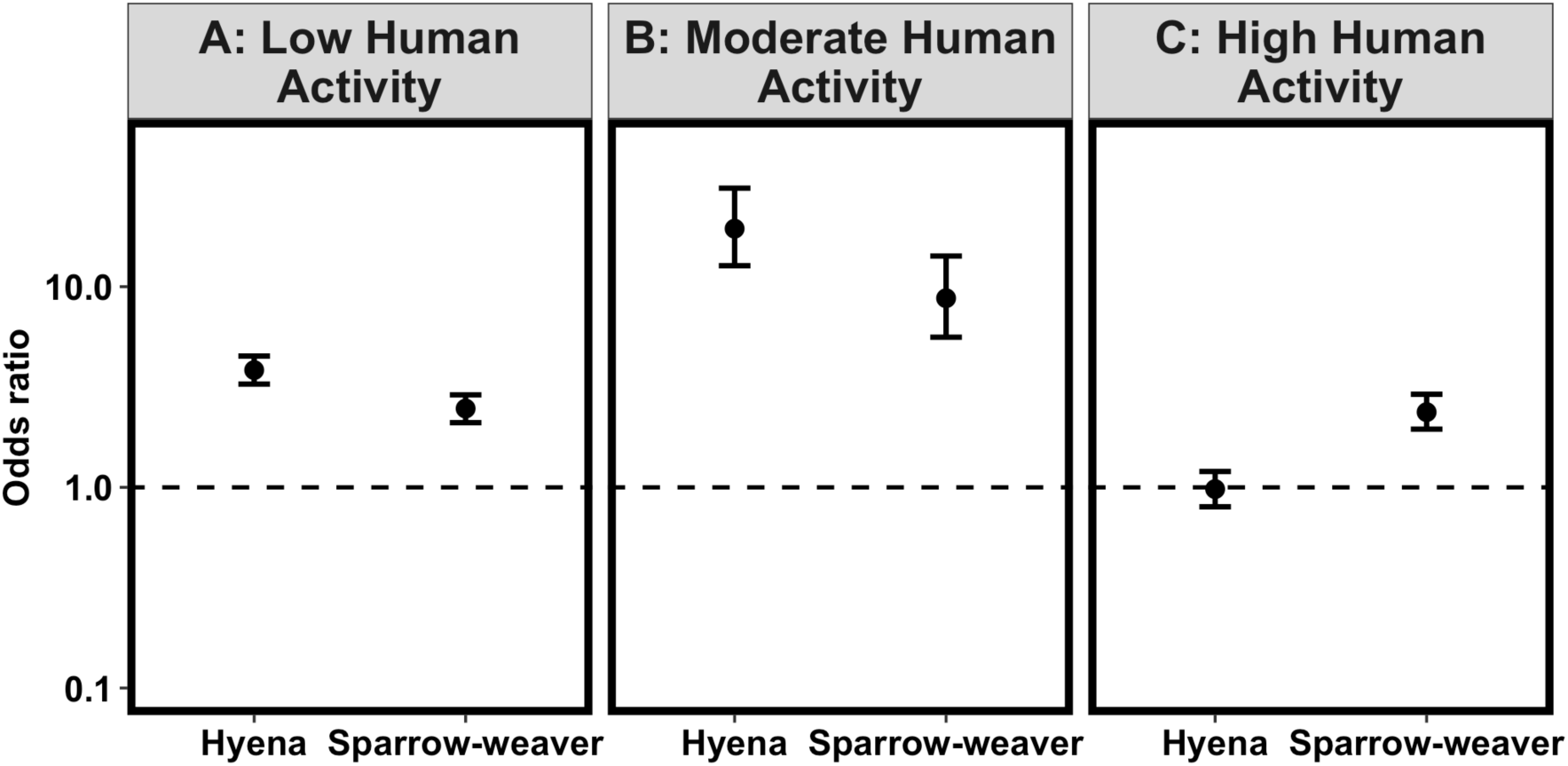
Odds ratios associated with dik-dik responses to different types of predation cues across the human activity gradient: areas of low (A), moderate (B) and high (C) human activity. Across each area, dik-diks respond more strongly to both predator-specific predation cues (hyena vocalizations) and non-specific predation cues (sparrow-weaver alarm calls) compared to the control treatment (sooty boubou song, depicted by the dashed line). However, the response of dik-diks to predator-specific predation cues in areas of high activity (C) did not differ significantly from their response to the control treatment. Error bars represent 95% confidence intervals.

Dik-diks were less vigilant and less flighty with increasing human activity. Over the 80-second recording periods, on average, dik-diks spent 20 ± 6 seconds of the monitoring period in a state of vigilance and 4 ± 4 seconds fleeing in areas of low human activity when responding to sooty boubou song. In areas of moderate human activity, the odds of dik-diks responding decreased by 85% (95% CI: -90 – -77%; *p* < 0.001), and dik-diks spent 5 ± 3 seconds of the monitoring period in a state of vigilance but never fled. In areas of high human activity, the odds of dik-diks responding decreased by 57% (95% CI: -65 – -44%; *p* < 0.001), and dik-diks spent 12 ± 8 seconds of the monitoring period in a state of vigilance but never fled.

The strength of response to hyena vocalizations decreased with increasing human activity (Fig. 4B). In areas of low human activity, dik-diks spent 17 ± 6 seconds of the monitoring period in a state of vigilance and 33 ± 9 seconds fleeing when responding to hyena vocalizations. In areas of moderate human activity, the odds of dik-diks responding decreased by 22% (95% CI: -37 – -3%; *p* < 0.001), and dik-diks spent 33 ± 3 seconds of the monitoring period in a state of vigilance and 13 ± 7 seconds fleeing. In areas of high human activity, the odds of dik-diks responding decreased by 89% (95% CI: -91 – -87%; *p* < 0.001), and dik-diks spent only 9 ± 3 seconds of the monitoring period in a state of vigilance and 3 ± 2 seconds fleeing.

**Figure 4:**
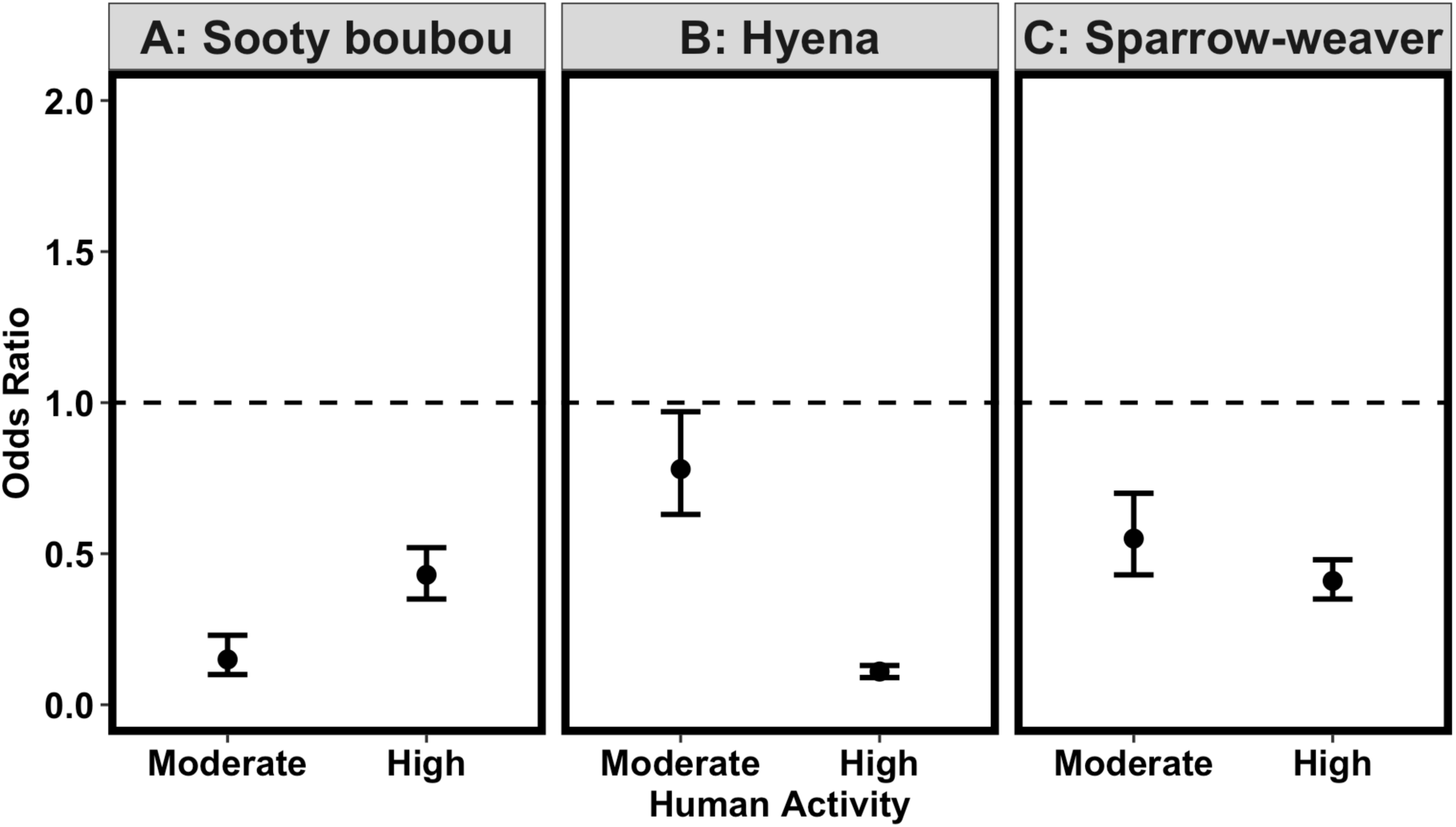
Odds ratios associated with dik-dik responses to different types of predation cue: control (sooty boubou song; A), predator-specific (hyena vocalizations; B) and non-specific (sparrow-weaver alarm calls; C). Except for the control treatment, the strength of response by dik-diks to predation cues decreased with increasing human presence. Error bars represent 95% confidence intervals, and the dashed line represents a response identical to the reference category (low human activity).

The strength of responses to sparrow-weaver alarm calls also decreased with increasing human activity (Fig. 4C). In areas of low human activity, dik-diks spent 39 ± 7 seconds of the monitoring period in a state of vigilance and 2 ± 1 seconds fleeing when responding to sparrow-weaver alarm calls. In areas of moderate human activity, the odds of dik-diks responding decreased by 45% (95% CI: -57 – -30%; *p* < 0.001), and dik-diks spent 30 ± 16 seconds of the monitoring period in a state of vigilance but never fled. In areas of high human activity, the odds of dik-diks responding decreased by 59% (95% CI: -65 – -52; *p* < 0.001) and dik-diks spent 16 ± 4 seconds of the monitoring period in a state of vigilance and 9 ± 5 seconds fleeing.

## 4. Discussion

We hypothesized that dik-diks in areas of higher human activity would respond less strongly to humans (Hypothesis 1), predator-specific predation cues (Hypothesis 2), and non-specific predation cues (Hypothesis 3) than dik-diks living in areas of lower human activity. Further, we hypothesized that dik-diks in areas of high human activity would respond more strongly to non-specific predator cues than predator-specific cues (Hypothesis 4). All four hypotheses were supported. Flight initiation distances and responses to recorded spotted hyena vocalizations, white-browed sparrow-weaver alarm calls, and sooty boubou song (simulating a predator-specific cue of predation risk, a non-specific cue of predation risk, and a control treatment, respectively) decreased sequentially as human activity increased, indicating that dik-diks have habituated to humans where there is more human activity (Hypothesis 1; Fig. 2). Similarly, dik-diks responded more strongly to both predator-specific and non-specific predation cues than controls, and the strength of responses to predator-specific (Hypothesis 2; Fig. 4B) and non-specific (Hypothesis 3; Fig. 4C) predation cues decreased with increasing human activity. Dik-diks, however, did not respond more strongly to predator-specific cues than the control treatment in areas of high human activity (Hypothesis 4; Fig. 3C). Overall, our results indicate that dik-diks exposed to higher levels of human activity are progressively more habituated to humans and naive to predation cues from non-human predators, but that naivety applies only to predators (e.g., spotted hyenas) that are excluded by human activity.

Habituation to humans varied with exposure to different levels of human activity (Hypothesis 1; Fig. 2). Upon detection of predator cues, flight improves fitness by avoiding subsequent pursuit by predators (Samia et al., 2013; Williams et al., 2014). However, as a response to benign interactions with humans, flight is maladaptive and can come at the expense of feeding and other behaviors that promote fitness (Cooper et al., 2015). Habituation is widespread across taxa including Nubian ibex (*Capra nubiana*; Saltz et al., 2019), rock and bush hyraxes (*Procavia capensis johnstoni* and *Heterohyrax brucei*; Mbise et al., 2020), inland blue-tailed skinks (*Emoia impar*; McGowan et al., 2014), and brush-turkeys (*Alectura lathami*; Hall et al., 2020).

Because habituation to humans allows prey to exploit human shields, habituation has knock-on effects for antipredator behaviors (Bar-Ziv et al., 2023). First, habituation to human activity was associated with weaker responses by dik-diks to predation cues (Hypothesis 2; Fig. 4B and C). This suggests that increased human activity and resulting naivety confers tolerance of non-human predators as well, which could be detrimental to prey populations if predators recolonize a human-occupied area. Because of the tendency for predators to avoid humans, behavior of prey in areas of high human activity can become decoupled from cues that prey would otherwise use to inform risk of predation. In our study, dik-diks’ weaker responses to predator-specific predation cues in areas of high human activity suggests predator naivety by dik-diks. As human populations expand and increasingly come to coexist with large predators, increased human activity may increase susceptibility of dik-diks and other wildlife to their non-human predators.

Given that habituation to humans can lead to relaxed antipredator behavior by prey, relatively benign but frequent interaction with humans such as ecotourism may have unintended consequences for prey species. This is particularly true when such activity occurs on multiple-use landscapes such as working ranches where predators maintain fear of humans. Since ecotourism is among the leading agents of economic growth in Sub-Sahara Africa (Nene & Taivan, 2017), and because ecotourism results in behavioral alteration by prey species (Geffroy et al., 2015; Saltz et al., 2019; Barocas et al., 2022), this may increase susceptibility of prey species to predation if, for example, a working ranch commits fully to ecotourism. As a caveat, however, and because dik-diks exhibit more antipredator behavior in areas with moderate levels of human activity than in areas of high human activity (Fig. 3B and C respectively), intermittent tourist activities such as safari excursions with temporary bouts of camping may impact prey’s ability to respond to predation cues less than in areas with permanent human activity. Because of the intermittent nature of many tourist activities, future research that aims to identify thresholds of tourist activity that lead to habituation and subsequent loss of antipredator behavior will be useful for the conservation of prey species.

Our findings contradict those of Coleman et al. (2008), who found in the same study system that habituated dik-diks were more able than non-habituated dik-diks to discriminate between a nonthreatening songbird and a predator-specific predation cue. This may be because Coleman et al. (2008) used distance to human settlement as a proxy for habituation, but did not experimentally measure levels of habituation using flight initiation distance to determine a threshold between their high- and low-habituation categories (Gil & Brumm, 2013). It is therefore possible that some “high-habituation” dik-diks in their study were relatively unhabituated to humans, while some “low-habituation” dik-diks were somewhat habituated to humans. Further, fear of humans tends to repel large predators more strongly than mesopredators (Suraci et al., 2019). Coleman et al. (2008) used jackal vocalizations as their predation cue, but jackals may not avoid human activity as strongly as larger predators like hyenas (and may even be attracted if food subsidies are present in areas of high human activity). This could explain why habituated dik-diks in Coleman et al. (2008) responded to jackal calls. Future research that accounts for human habituation as a continuous predictor and that includes a wider variety of predator vocalizations may reconcile our results with those of Coleman et al. (2008). Nevertheless, because Coleman et al. (2008) found that habituation does not affect the ability of prey species to perceive risk of predation, they suggested that habituation may not affect survival of captive-bred, translocated, or reintroduced animals. In contrast, we found that habituation leads to relaxed antipredator behavior among prey species, making them more vulnerable to predation. Our findings indicate that decreased predator awareness of captive-bred, translocated, or reintroduced animals that are habituated to humans may indeed reduce survival and fitness (Carthey & Blumstein, 2018).

Habituation to humans also resulted in reduced responses to non-specific predation cues (Hypothesis 3; Fig. 4C). Dik-diks eavesdrop on cues from unrelated species (such as white-bellied go-away birds; Lea et al., 2008) to avoid predation. Such heterospecific eavesdropping is common in two scenarios: (1) between two prey species that share a common predator (e.g., impala [*Aepyceros melampus*] and chacma baboons [*Papio hamadryas ursinus*] in Botswana; Kitchen et al., 2010), or (2) between a prey species and another species that is associated with the presence of a predator (e.g., moose [*Alces alces*] and common ravens [*Corvus corax*], which feed on carrion, implying that a predator has been hunting nearby; Berger, 1999). While our study focused on the relationship between two prey species that share common predators, investigations on whether prey species in areas with high human activity respond differently to cues from species that are simply associated with predators—such as scavenger vocalizations—will be invaluable for further understanding variation in heterospecific eavesdropping. In addition, studies focused on heterospecific eavesdropping (including our own) typically measure it as a dichotomous response—either it happens, or it does not—without accounting for variation in the degree of response between different prey populations and species. Future studies that consider how heterospecific eavesdropping changes with prey size would improve our knowledge of this phenomenon. For example, in our study area, larger prey that are not vulnerable to as many predators as dik-diks and therefore do not fear mesopredators may cease to eavesdrop on other species as they become habituated.

Interestingly, in areas of high human activity, the probability of dik-diks responding to non-specific predation cues was significantly higher than the probability of dik-diks responding to predator-specific predation cues (Fig. 3C). While hyena calls convey information only about the presence of hyenas (which strongly avoid humans, particularly during the day), the non-specific predation cue—the sparrow-weaver alarm call—could signal the presence of other dik-dik predators that do not avoid humans as strongly, such as snakes, raptors, or baboons (Kingswood & Kumamoto, 1996). Habituated dik-diks may only lose their sensitivity to hyenas and other large predators that avoid areas with high human activity, while remaining vigilant toward smaller predators that can remain prevalent even in areas of high human activity. Human activity tends to repel large predators more than mesopredators, which may be attracted to human activity to avoid predation by large predators and exploit food subsidies (Suraci et al., 2016, 2019). Our findings indicate that dik-diks can distinguish between different types of predation cue, and that they can titrate their levels of response accordingly. This could further explain why habituated dik-diks respond to vocalizations from jackals (a mesopredator; Coleman et al. 2008) but did not respond to vocalizations from hyenas (a large predator) in our study. Additional research that tests responses of prey species to different classes of predators (e.g., mesopredators vs. apex predators, ambush vs. cursorial predators) in areas with varying levels of human activity would be useful for further characterizing the extent to which habituation to humans alters antipredator behavior.

## Conclusions

Dik-diks living in areas of high human activity were more habituated and spent less time responding to predation cues than those in areas with less human activity. As human populations expand and increasingly co-occur with large predators, increased human activity may increase susceptibility of dik-diks and other wildlife to their non-human predators. These results are particularly relevant for ecotourism in working landscapes and translocation or reintroduction of habituated animals.

## Acknowledgements

We would like to thank the Tropical Biology Association (TBA) for organizing the field course and giving us opportunity to learn how to better conduct field research. Many thanks to all the tutors of the 2022 field course, especially Rosie Trevelyan, Duncan Kimuyu, Jeff Ollerton and Kimani Ndung’u, and to all the students for sharing their knowledge. A special thanks to Rosie Trevelyan for ideas during our planning stage; Kim Mortega for teaching us how to record animals and edit sound using SASlab Pro, and better conduct behavioral studies; Maximillian Schotts and Emma (Kim’s husband and daughter respectively) for helping spot dik-diks during our study. A special thanks to all the TBA support staff, especially James Gichia for driving us, and Mariana Carvalho for planning to ensure every group had resources to conduct fieldwork. We would also like to thank Mpala Research Centre for hosting us, their support staff for their hospitality, and rangers for protection against buffalo and elephants while conducting our fieldwork. Funding for this research was provided by the Conservation Leadership Programme and the School of Natural Resources and the Environment at the University of Arizona.

