## Supplementary Material for "Human shields alter antipredator behavior in Guenther’s dik-dik (*Madoqua guentheri*)"

### *Summary*

This Supplementary Material provides the results on how human activity affected the proportion of time spent by dik-diks in response to predation cues using beta regression analysis. Besides response to sparrow-weaver alarm call (Fig S1, A), the results are consistent with our analysis using binomial-family generalized linear models: where there are more human, we expect dik-dik to have a decreased response to vocalization from hyena but retain some response to sparrow-weavers. With this analysis however, the confidence intervals, and *p-values* are wider and more conservative respectively because beta regression estimates more parameters than generalized linear models.

### *Results*

In areas of low human activity (Fig S1, A), the odds of dik-diks responding increased by 170 % (95% CI: 116 – 630%; $p$ = 0.021) to hyena vocalizations but did not differ (95% CI: 88 – 401%; $p$ = 0.10), when responding to sparrow-weaver, compared to the sooty boubou song. In areas of moderate human activity (Fig. S1, B), the odds of dik-diks responding increased by 408% (95% CI: 58–4456%; $p$ = 0.14) and 251% (95% CI: 87–1445%; $p$ = 0.078) when responding to hyena vocalizations and sparrow-weaver alarm calls, respectively, compared to the sooty boubou song. In areas of high human activity (Fig. S1, C), the odds of dik-diks responding did not differ (95% CI: 54–587%; $p$ = 0.35) when responding to hyena vocalizations compared to sooty boubou song, but increased by 198% (95% CI: 0.87–9.97; $p$ = 0.082) when responding to and sparrow-weaver alarm calls, compared to the sooty boubou song.

When responding to sooty boubou, in areas of moderate human activity (Fig S2, A), the odds of dik-diks responding decreased by 65% (95% CI: -92.00 – 165%; $p$ = 0.19), and dik-diks spent 5 ± 3 seconds of the monitoring period in a state of vigilance but never fled. In areas of high human activity, the odds of dik-diks responding decreased by 65% (95% CI: -90 – 126%; $p$ = 0.11) and dik-diks spent 12 ± 8 seconds of the monitoring period in a state of vigilance but never fled.

The strength of response to hyena vocalizations decreased with increasing human activity (Fig. S2, B). In areas of low human activity, dik-diks spent 17 ± 6 seconds of the monitoring period in a state of vigilance and 33 ± 9 seconds fleeing when responding to hyena vocalizations. In areas of moderate human activity, the odds of dik-diks responding decreased by 24% (95% CI: - 71.00 – 196%; $p$ = 0.56), and dik-diks spent 33 ± 3 seconds of the monitoring period in a state of vigilance and 13 ± 7 seconds fleeing. In areas of high human activity, the odds of dik-diks responding decreased by 81% (95% CI: -93.00 – -43.00; $p$ = 0.0007), and dik-diks spent only 9 ± 3 seconds of the monitoring period in a state of vigilance and 3 ± 2 seconds fleeing.

The strength of responses to sparrow-weaver alarm calls also decreased with increasing human activity (Fig. S2. C). In areas of low human activity, dik-diks spent 39 ± 7 seconds of the monitoring period in a state of vigilance and 2 ± 1 seconds fleeing when responding to sparrow-weaver alarm calls. In areas of moderate human activity, the odds of dik-diks responding decreased 47% (95% CI: -87.00 – 114.00%; $p$ = 0.38), and dik-diks spent 30 ± 16 seconds of the monitoring period in a state of vigilance but never fled. In areas of high human activity, the odds of dik-diks responding decreased by 58% (CI: -83.00 – 2.00; $p = 0.054$), and dik-diks spent 16 ± 4 seconds of the monitoring period in a state of vigilance and 9 ± 5 seconds fleeing.

*Plot outputs*

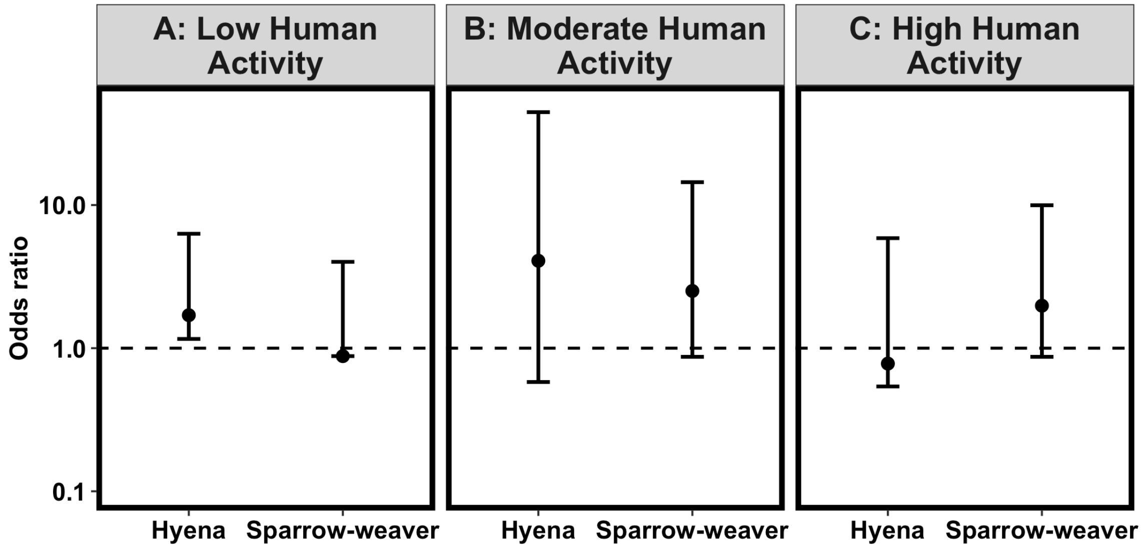


*Figure S1: The odds ratio associated with dik-dik responses to different types of predation cues across the human activity gradient: areas of low (A), moderate (B) and high (C) human activity. Across each area, dik-diks respond more strongly to both predator-specific predation cues (hyena vocalizations) and non-specific predation cues (sparrow-weaver alarm calls) compared to the control treatment (sooty boubou song, depicted by the dashed line). However, the response of dik-diks to non-specific predation cues and predator-specific predation cues in areas of low (A) and high (C) activity respecitvely did not differ significantly from their response to the control treatment. Error bars represent 95% confidence intervals.*

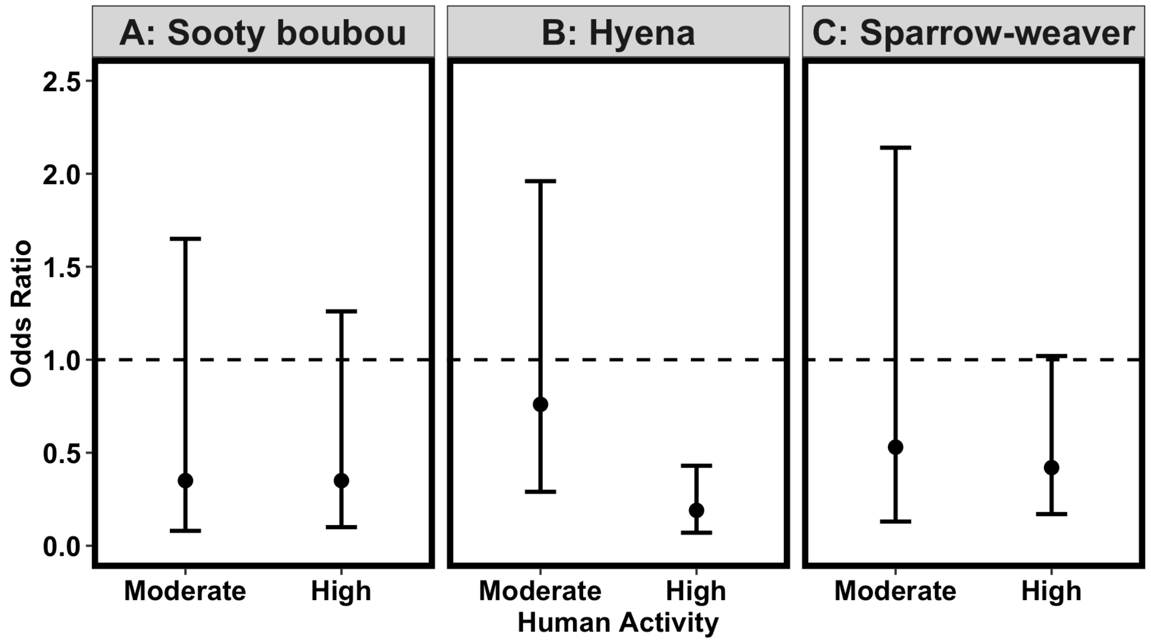


*Figure S2: Odds ratios associated with dik-dik responses to different types of predation cue: control (sooty boubou song; A), predator-specific (hyena vocalizations; B) and non-specific (sparrow-weaver alarm calls; C). Except for the control treatment, the strength of response by dik-diks to predation cues decreased with increasing human presence. Error bars represent 95% confidence intervals, and the dashed line represents a response identical to the reference category (low human activity).*
